# Characterising undiagnosed chronic obstructive pulmonary disease: a systematic review and meta-analysis

**DOI:** 10.1101/184986

**Authors:** Kate M. Johnson, Stirling Bryan, Shahzad Ghanbarian, Don D. Sin, Mohsen Sadatsafavi

## Abstract

**Background:** A significant proportion of patients with chronic obstructive pulmonary disease (COPD) remain undiagnosed. Characterising these patients can increase our understanding of the ‘hidden’ burden of COPD and the effectiveness of case detection interventions.

**Methods:** We conducted a systematic review and meta-analysis to compare patient and disease risk factors between patients with undiagnosed persistent airflow limitation and those with diagnosed COPD. We searched MEDLINE and EMBASE for observational studies of adult patients meeting accepted spirometric definitions of COPD. We extracted and pooled summary data on the proportion or mean of each risk factor among diagnosed and undiagnosed patients (unadjusted analysis), and coefficients for the adjusted association between risk factors and diagnosis status (adjusted analysis). This protocol is registered with PROSPERO (CRD42017058235).

**Findings:** 2,083 records were identified through database searching and 16 articles were used in the meta-analyses. Diagnosed patients were less likely to have mild (v. moderate to very severe) COPD (odds ratio [OR] 0·30, 95% CI 0·24-0·37, 6 studies) in unadjusted analysis. This association remained significant but its strength was attenuated in the adjusted analysis (OR 0·72, 95% CI 0·58-0·89, 2 studies). Diagnosed patients were more likely to report respiratory symptoms such as wheezing (OR 3·51, 95% CI 2·19-5·63, 3 studies) and phlegm (OR 2·16, 95% CI 1·38-3·38, 3 studies), had more severe dyspnoea (modified Medical Research Council scale mean difference 0·52, 95% CI 0·40-0·64, 3 studies) and slightly greater smoking history than undiagnosed patients. Patient age, sex, current smoking status, and the presence of coughing were not associated with a previous diagnosis.

**Interpretation:** Patients with undiagnosed persistent airflow limitation had less severe airflow obstruction and fewer respiratory symptoms than diagnosed patients. This indicates that there is lower disease burden among undiagnosed patients compared to those with diagnosed COPD, which may significantly delay the diagnosis of COPD.

**Funding:** Canadian Institutes of Health Research.

**Declaration of interests:** We declare no competing interests.

**Author Contributions:** MS, SB, and KJ formulated the study idea and designed the study. KJ and SG performed all data analyses and MS, SB and DS contributed to interpretation of findings. KJ wrote the first draft of the manuscript. All authors critically commented on the manuscript and approved the final version. MS is the guarantor of the manuscript.

## Research in context

### Evidence before this study

Many cross-sectional prevalence studies have compared the characteristics of patients with persistent airflow limitation but no prior diagnosis of COPD (‘undiagnosed’) to those with persistent airflow limitation and a diagnosis of COPD (‘diagnosed’). We searched MEDLINE and EMBASE for observational studies published in English between January 1, 1980 and April 11, 2017 that assessed diagnosis status among adult patients with spirometrically defined persistent airflow limitation. We used search terms relating to COPD (including “chronic obstructive pulmonary disease” OR “bronchitis” OR “emphysema”) AND diagnosis (“diagnostic errors” OR “undiagnosed) AND risk factors (“risk factors” OR “characteristics”) to identify references. 18 articles met the eligibility criteria and 16 were included in the meta-analysis. Approximately half of the 18 eligible studies used random sampling of the general population; the other half used convenience sampling (e.g., recruitment from health-care settings) and tended to score lower in our qualitative quality assessment. We used summary data from the included articles to generate pooled estimates of the associations between sex, age, current smoking status, smoking history, respiratory symptoms, disease severity, and the likelihood of having received a previous diagnosis of COPD. Overall, disease characteristics had much greater discriminatory ability than patient characteristics, and more severe disease was the most characteristic of patients with ‘diagnosed’ COPD, followed by the presence of respiratory symptoms. Patients with ‘diagnosed’ COPD were 70% less likely to have mild disease compared to moderate, severe, or very severe COPD, and they were two to five times more likely to report the presence of respiratory symptoms such as wheeze, phlegm, and dyspnoea.

### Added value of this study

Lamprecht et al. found that undiagnosed patients tended to be younger male never smokers with fewer respiratory symptoms and less severe COPD using individual data from four population-based studies. Our study extends these findings by providing pooled estimates of the associations between patient and disease factors and the likelihood of receiving a diagnosis of COPD. Our results confirm the strong association between disease severity, respiratory symptoms, and COPD diagnosis that was previously reported. However, pooled estimates from 16 studies revealed no association between patient characteristics (age, sex) and COPD diagnosis, and only a weak association with smoking history. These estimates were consistent across alternate definitions of persistent airflow limitation, the population sampled (general population v. health-care setting), and analysis methods (contingency tables v. regression models). Our study provides a robust and generalizable characterisation of patients with undiagnosed persistent airflow limitation.

### Implications of all the available evidence

This systematic review and meta-analysis provides strong evidence that undiagnosed patients tend to have milder disease and fewer symptoms. Our findings can be used as selection criteria to target subgroups of patients with a high prevalence of underdiagnosis for case detection or screening. They also show that the burden of disease is lower in patients with undiagnosed persistent airflow limitation than in those with diagnosed COPD, indicating that there is a substantial lag between the development of persistent airflow limitation and receiving a diagnosis of COPD. This delay is a key missed opportunity to modify risk factors at the critical early stages of disease development.

## Introduction

Chronic Obstructive Pulmonary Disease (COPD) is an inflammatory lung disorder that is characterised by persistent airflow limitation^1^ and associated with symptoms of shortness of breath, cough and sputum production.^2^ Patients with COPD generally seek medical attention when they experience respiratory symptoms, most notably dyspnoea that is persistent and progressive.^1^ However, owing to under-utilization of lung function measurements and non-specific nature of the symptoms, COPD is often not recognized until late in the disease process. Indeed, many patients do not receive a diagnosis of COPD until after being hospitalized due to a severe exacerbation.^3^

Lamprecht et al.^4^ reported an average underdiagnosis rate of 81% in a prevalence study that included 30,874 participants across 44 countries. Reducing risk factors such as smoking and occupational risk factors while the disease is early in its progression is an important component of treatment for COPD.^5^ As such, late diagnosis of COPD represents a missed opportunity to modify the course of the disease through evidence-informed risk factor management and treatment.^6,7^ The extent of this missed opportunity is a function of both the number of COPD patients who are undiagnosed, as well as the burden of disease (e.g., symptom burden, lung function status) in this population.

Quantifying the true burden of undiagnosed COPD can be informed by a comparative assessment of patient-and disease-factors between diagnosed and undiagnosed patients. Numerous studies have compared the characteristics of patients with undiagnosed and diagnosed COPD, but to the best of our knowledge, these studies have never been systematically compiled and pooled. We hypothesized that the characteristics of patients, their risk factors, respiratory symptoms, and disease stage influence the likelihood of sreceiving a diagnosis of COPD.

## Methods

### Search strategy and selection criteria

We conducted a systematic review and meta-analysis to compare patient characteristics, risk factors, and symptoms in diagnosed and undiagnosed patients. We searched MEDLINE and EMBASE using the Ovid interface for eligible articles. The search strategy (Appendix, Text A1) was developed in MEDLINE and adapted to EMBASE using appropriate vocabulary terms. We included longitudinal or cross-sectional studies published in English between 1980 and April 11, 2017 that were based on original analysis of individual data. We did not include conference abstracts unless they met the inclusion criteria and provided the required information, and we did not assess grey literature. We extracted summary data from the eligible articles and contacted the authors to obtain additional information when required (one author group provided us with additional information). Title and abstract screening were initially performed, followed by full-text analysis to determine article eligibility. We extracted data using a customized Excel spreadsheet after the eligible articles had been compiled. KJ initially performed the selection procedure, and SG independently repeated each step on a subset (10%) of articles. Discrepancies were resolved through discussions between the two reviewers. Duplicate articles found in both MEDLINE and EMBASE were identified using a reference manager and manually removed. We used the Quality Assessment Tool for Observational Cohort and Cross-Sectional Studies developed by the National Institutes of Health National Heart, Lung, and Blood Institute^8^ to assign an overall quality rating (good, fair, or poor) to each study. KJ extracted relevant data and assessed the quality of the included studies, and SG replicated the assessment on 10% of articles. The reviewers determined the overall quality of each article by assigning ‘yes’, ‘no’, or ‘other’ (cannot determine, not applicable, or not reported) to 14 questions relating to external validity, bias in the measurements of the risk factors or outcomes, and confounders present in the study. The results of this assessment were assessed qualitatively.

The population of interest in this review were adult patients (≥18 years old) with persistent airflow limitation at the time of assessment. Persistent airflow limitation was defined when the study subjects demonstrated a ratio of Forced Expiratory Volume in 1 Second (FEV1) to Forced Vital Capacity (FVC) < 0·7 (fixed ratio definition)^1^ or FEV1 to FVC lower than the lower limit of normal (LLN definition)^9^ after the administration of a bronchodilator during spirometry. Study subjects who had airflow limitation and also a prior diagnosis of COPD or an obstructive lung disease (emphysema, chronic bronchitis, asthma) from a health-care professional were considered to have ‘diagnosed’ COPD, whereas those with persistent airflow limitation but without a prior health professional diagnosis of COPD were considered to be ‘undiagnosed’. Patients with other respiratory diseases were excluded. We included studies that sampled patients from any population or health-care setting.

Given the exploratory nature of the observational studies included in this review, we used a broad definition of risk factors that included any observable factor that could be associated with the probability of having received a diagnosis of COPD. Risk factors included patient-reported respiratory symptoms (cough, wheeze, phlegm, dyspnoea), sex, age, current smoking status, smoking history (pack-years), and disease severity classified using the Global Initiative for chronic Obstructive Lung Disease (GOLD) grades. The relationship of interest was the association between these risk factors and the probability of having ‘diagnosed’ COPD among patients with persistent airflow limitation.

We extracted summary data from each eligible article, which included study characteristics, the definition of persistent airflow limitation that was employed in each of the studies, the method of COPD diagnosis, and sample size. We also extracted the proportion or mean of risk factors between the diagnosed and undiagnosed groups, as well as the odds ratios (ORs) and their confidence intervals in studies that used regression modelling to assess the independent impact of the risk factors on diagnosis status. The protocol for this study is registered on the PROSPERO register of systematic reviews (CRD42017058235).^10^

### Data analysis

We used data extracted from articles measuring categorical data to generate ORs and standard errors for the association between risk factors and the probability of having received a diagnosis of COPD. In articles assessing continuous data, we calculated the mean difference (MD) in risk factors and their standard errors among diagnosed and undiagnosed patients. We pooled the ORs or MDs from individual studies using the inverse variance method implemented with the ‘meta’ package^11^ in R Statistical Software^12^ (version 3.3.3). We used fixed-effects models when estimates from only two studies were being pooled, or if the null hypothesis that all studies evaluated the same effect was not rejected (at 0·05 significance level) using Cochran’s Q statistic.^13^

Otherwise, we used random-effects models. We quantified heterogeneity between studies 2 14 using the I statistic. We did not pool together studies that used alternate definitions of persistent airflow limitation (fixed ratio and LLN). When separate studies used subsets of the same dataset (i.e., the Latin American Project for the Investigation of Obstructive Lung Disease [PLATINO] dataset^4,15–17^), we used the estimate from the study with the largest sample size. We conducted a sensitivity analysis to determine the association between the risk factors and COPD diagnosis only among population-based studies (those based on random sampling of the general population as opposed to convenience sampling).

## Results

The search resulted in 1,857 references after excluding duplicates. 1,788 references were excluded by screening their titles and abstracts, and 69 remained for full text review to determine eligibility. A total of 18 articles met the inclusion criteria following the screening process, but only 16 articles were included in quantitative synthesis (Figure 1). The overall agreement between reviewers was high (90%).

**Figure 1:**
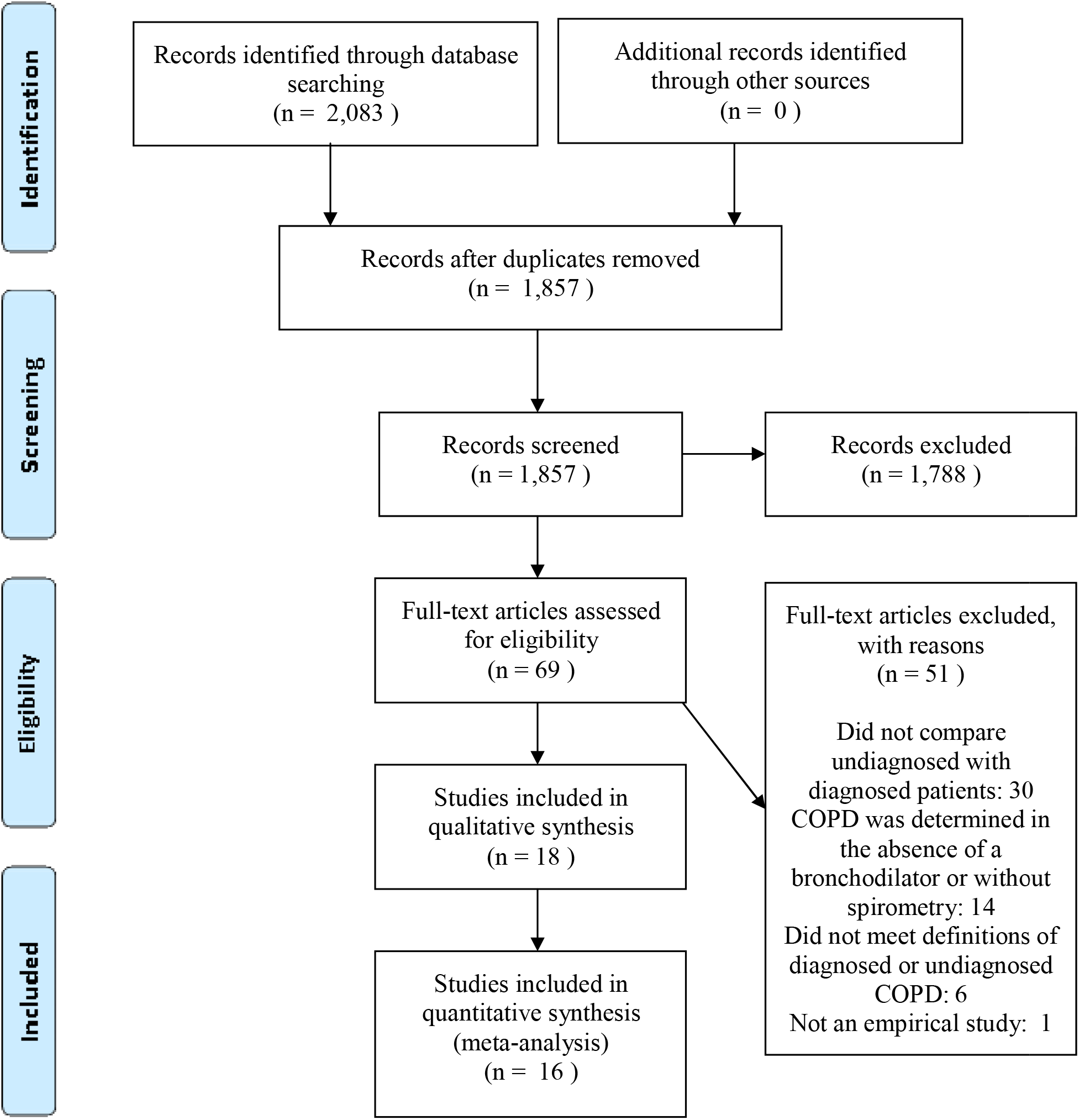
Preferred Reporting Items for Systematic Reviews and Meta-Analyses flow diagram.

A summary of the 18 eligible articles is presented in Table 1. However, two eligible articles were excluded from the meta-analysis because they were missing the necessary information,^18^ or did not measure any risk factors in common with other studies.^19^ The majority of the 18 eligible articles were cross-sectional (n=16), and were population-based (n=10). Other studies sampled patients from primary care clinics (n=4), hospitalized patients (n=3), or participants in a smoking cessation program (n=1). Studies originated from Latin America (n=6), Europe (n=6), Canada (n=2), and Asia (n=2). Data from the Epidemiologic Study of COPD in Spain (EPI-SCAN),^4,20,21^ PLATINO, and the Burden of Obstructive Lung Disease (BOLD),^4,22^ were used in three, four, and two different studies, respectively, but only one study from each dataset was included in pooled analyses. The definition of persistent airflow limitation varied between articles; 15 studies defined it as the fixed ratio, two studies used the LLN definition, and one study reported results using both definitions. The percentage of patients with undiagnosed persistent airflow limitation was greater than 50% in all but two studies (which sampled from health-care settings).

**Table 1:**
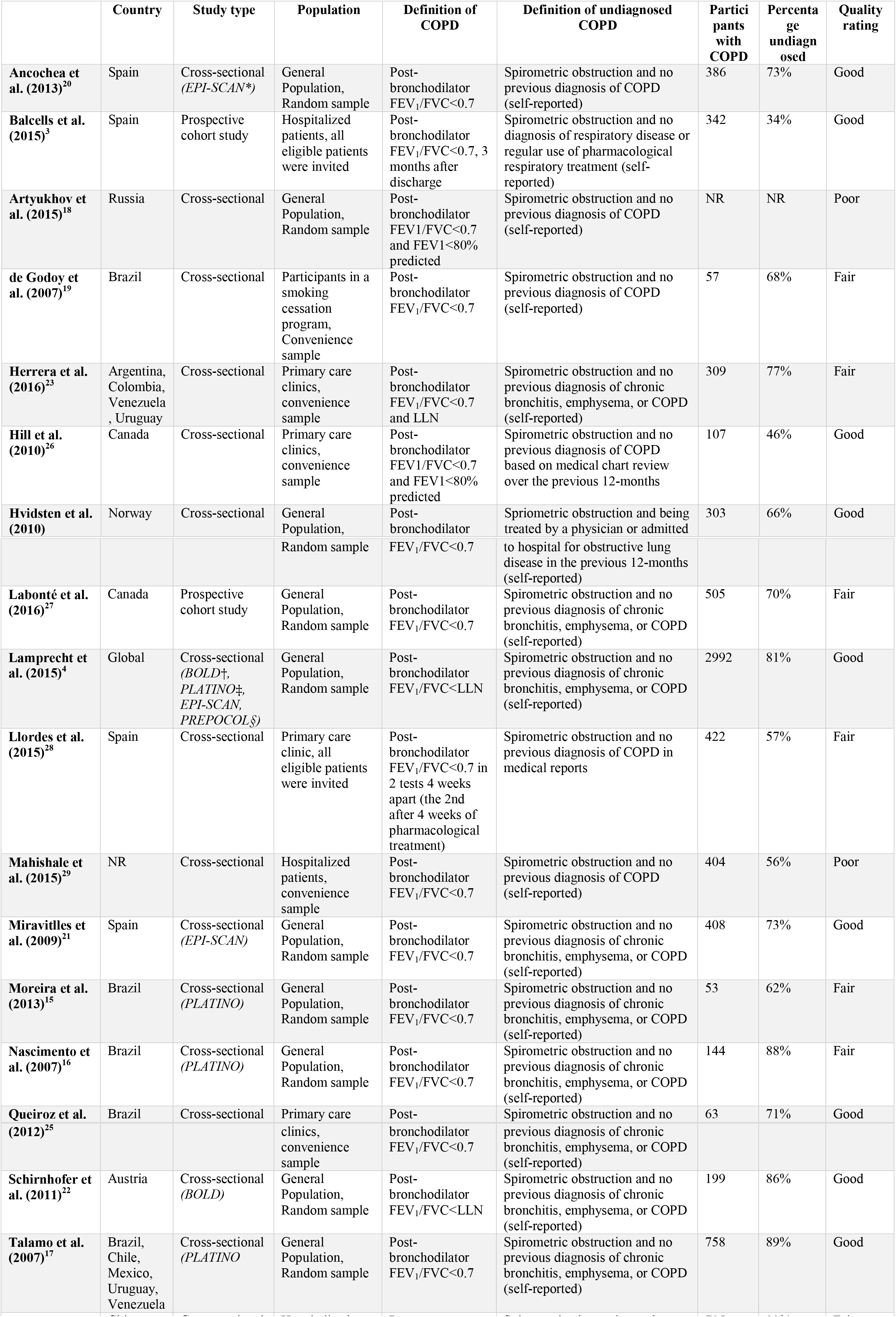
Characteristics of selected studies

Not Reported (NR)

^*^Epidemiologic Study of COPD in Spain (EPI-SCAN)

^†^Burden of Obstructive Lung Disease (BOLD)

^‡^Latin American Project for the Investigation of Obstructive Lung Disease (PLATINO)

^§^Prevalence study of COPD in Colombia (PREPOCOL)

The quality of the 18 eligible articles was variable. Half of the studies were assigned a quality rating of ‘good’, seven studies were assigned a rating of ‘fair’, and two studies were deemed poor in quality. Studies that were not assigned a ‘good’ quality rating generally had a primary study focus that was not our question of interest. For example, comparing the characteristics of diagnosed and undiagnosed patients was only reported tangentially in five studies, and disease severity was the only factor that was compared between the undiagnosed and diagnosed groups in three of the studies. The use of regression modelling to examine the independent impact of risk factors on the likelihood of receiving a COPD diagnosis was uncommon (performed in only seven studies), and in studies that used regression modelling, the risk factors that were adjusted for varied substantially.

### Unadjusted analysis

Comparisons of the characteristics of diagnosed and undiagnosed patients with persistent airflow limitation based on contingency tables (‘unadjusted analysis’) were reported in 12 studies. Because of the predominance of the fixed-ratio definition of airflow limitation, pooled results from studies that used this definition are reported in the main text and LLN-based results are provided in the Appendix. Pooled comparisons of sex, respiratory symptoms, current smoking status, smoking history, and COPD severity among patients meeting the fixed ratio definition of airflow limitation are shown in Figure 2.

**Figure 2:**
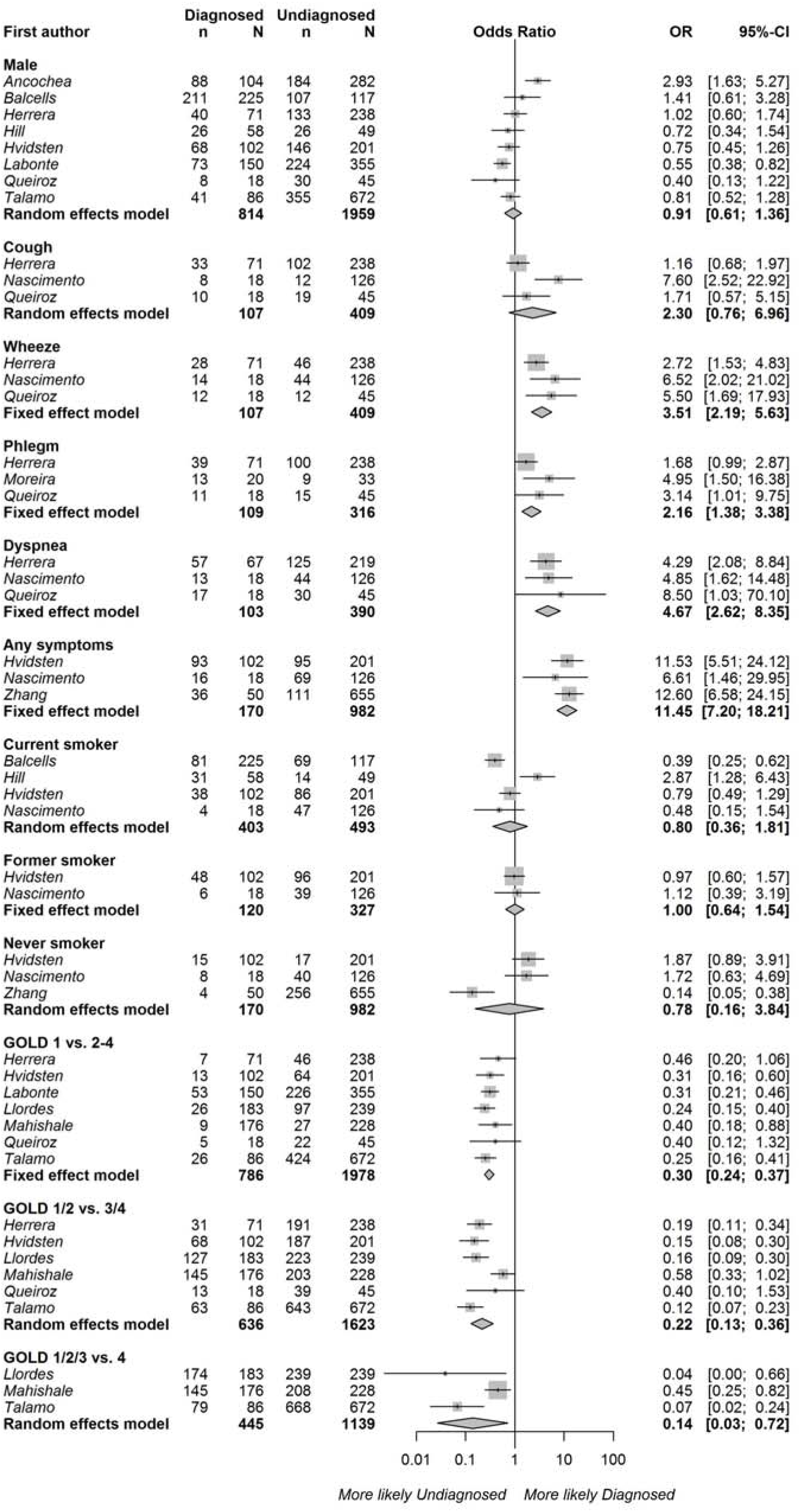
Associations between diagnosed (v. ‘undiagnosed’) COPD and sex, the presence of cough, wheeze, phlegm, dyspnoea, any respiratory symptoms, smoking status, smoking history, and COPD severity based on contingency tables. Persistent airflow limitation was defined as post-bronchodilator FEV1/FVC<0·7. Squares represent individual study estimates with the size of the square corresponding to their weight in the pooled estimate (represented with diamonds).

**Figure 3:**
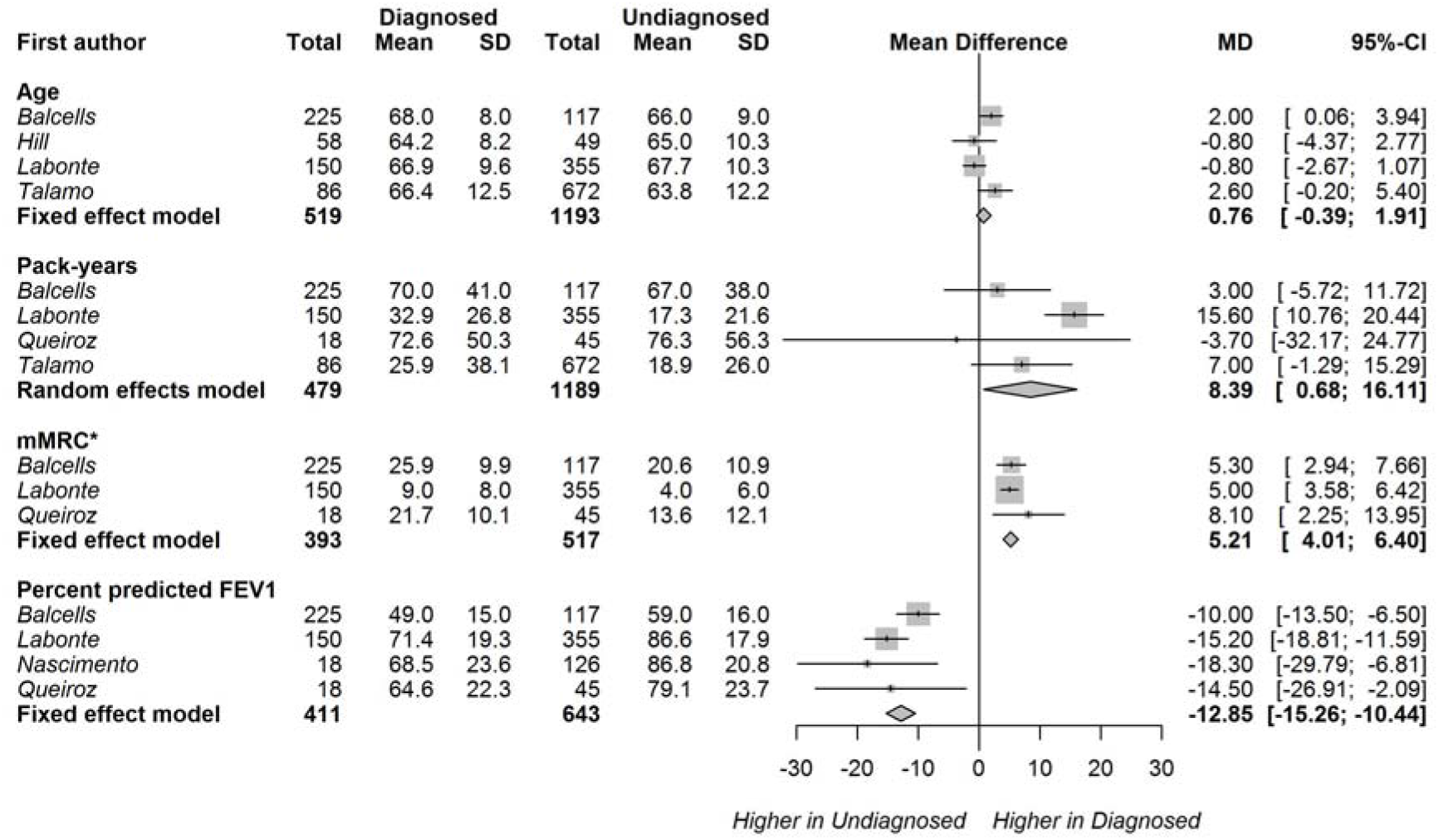
Mean difference (MD) in age, pack-years of smoking, mMRC dyspnoea score, and percent of predicted FEV1 between diagnosed and undiagnosed categories. Persistent airflow limitation was defined as post-bronchodilator FEV1/FVC<0·7. Squares represent individual study estimates with the size of the square corresponding to their weight in the pooled estimate (represented with diamonds). *modified Medical Research Council (mMRC) Dyspnoea scale^24^ means and standard errors (SE) for the diagnosed and undiagnosed categories are multiplied by a factor of 10.

Patients with ‘diagnosed’ COPD were more likely to be experiencing respiratory symptoms such as wheezing (OR 3·51, 95% CI 2·19-5·63, 3 studies), phlegm (OR 2·16, 95% CI 1·38-3·38, 3 studies), dyspnoea (OR 4·67, 95% CI 2·62-8·35, 3 studies), or any respiratory symptoms (OR 11·45 95% CI 7·20-18·21, 3 studies). They were much less likely to have mild (grade I) COPD than moderate to very severe COPD (grade II-IV) as measured by GOLD grades (OR 0·30 95% CI 0·24-0·37, 7 studies). The heterogeneity between studies was relatively low (I^2^<35.0% for wheeze, phlegm, dyspnoea, any_2_ symptoms, and COPD severity); however, the I statistics should be interpreted cautiously due to the low number of studies within each category. Patient sex, current smoking status, and smoking history were not associated with ‘diagnosed’ COPD. Having a cough was also not significantly associated with diagnosis status, however variability between the three studies measuring this risk factor was particularly high (I^2^77·9%).

Sensitivity analysis of only the population-based studies revealed very similar results (n=5 studies, Appendix, Figure A1). Pooled analysis of two studies^22,23^ using the LLN definition of airflow limitation was consistent with the findings based on fixed ratio results (Appendix, Figure A2); however, cough was marginally associated with diagnosis status in this analysis (OR 1·65, 95% CI 1·02-2·66).

Similarly, patients with ‘diagnosed’ COPD (fixed ratio definition) were more impaired by dyspnoea (modified Medical Research Council [mMRC] dyspnoea scale^24^ MD 0·52, 95% CI 0·40-0·64, 3 studies) and had greater airflow obstruction (percent predicted FEV1 MD −12·85%, 95% CI −15·26% to −10·44%, 4 studies) than undiagnosed patients. Patients with ‘diagnosed’ COPD also had a slightly greater smoking history (pack-years MD 8·39, 95% CI 0·68-16·44, 4 studies); however there was high variability between the study_2_ means (I 84·2%). There was no difference in mean age between diagnosed and undiagnosed patients.

### Adjusted analysis

Articles using regression modelling to assess the independent impact of risk factors on COPD diagnosis (‘adjusted analysis’) were pooled by risk factor type, and the results are presented in Figure 4 for the fixed ratio definition of persistent airflow limitation (5 articles), and Figure 5 for the LLN definition (2 articles with 5 datasets). The effect sizes of the risk factors were attenuated in these adjusted analyses. The presence of phlegm had a weak independent association with the diagnosis of COPD (OR 1·16, 95% CI 1·00-1·35, 2 studies) using the fixed ratio definition. The presence of wheezing (OR 1·20, 95% CI 0·99-1·44, 2 studies) and dyspnoea (OR 1·13 95% CI 0·99-1·29, 2 studies) were not independently associated with a diagnosis. In contrast, mild COPD (GOLD grade I OR 0·72, 95% CI 0·58-0·80) or moderate COPD (GOLD grade II, OR 0·71, 95% CI 0·58-0·86), were independently associated with a lower likelihood of diagnosis, compared with severe or very severe (reference GOLD grades III-IV). Sex and the presence of cough did not influence the likelihood of being diagnosed in the adjusted analyses, although the number of studies were small and heterogeneity in the effect estimates between studies was very high (I^2^>70.0% for all risk factors except sex).

**Figure 4:**
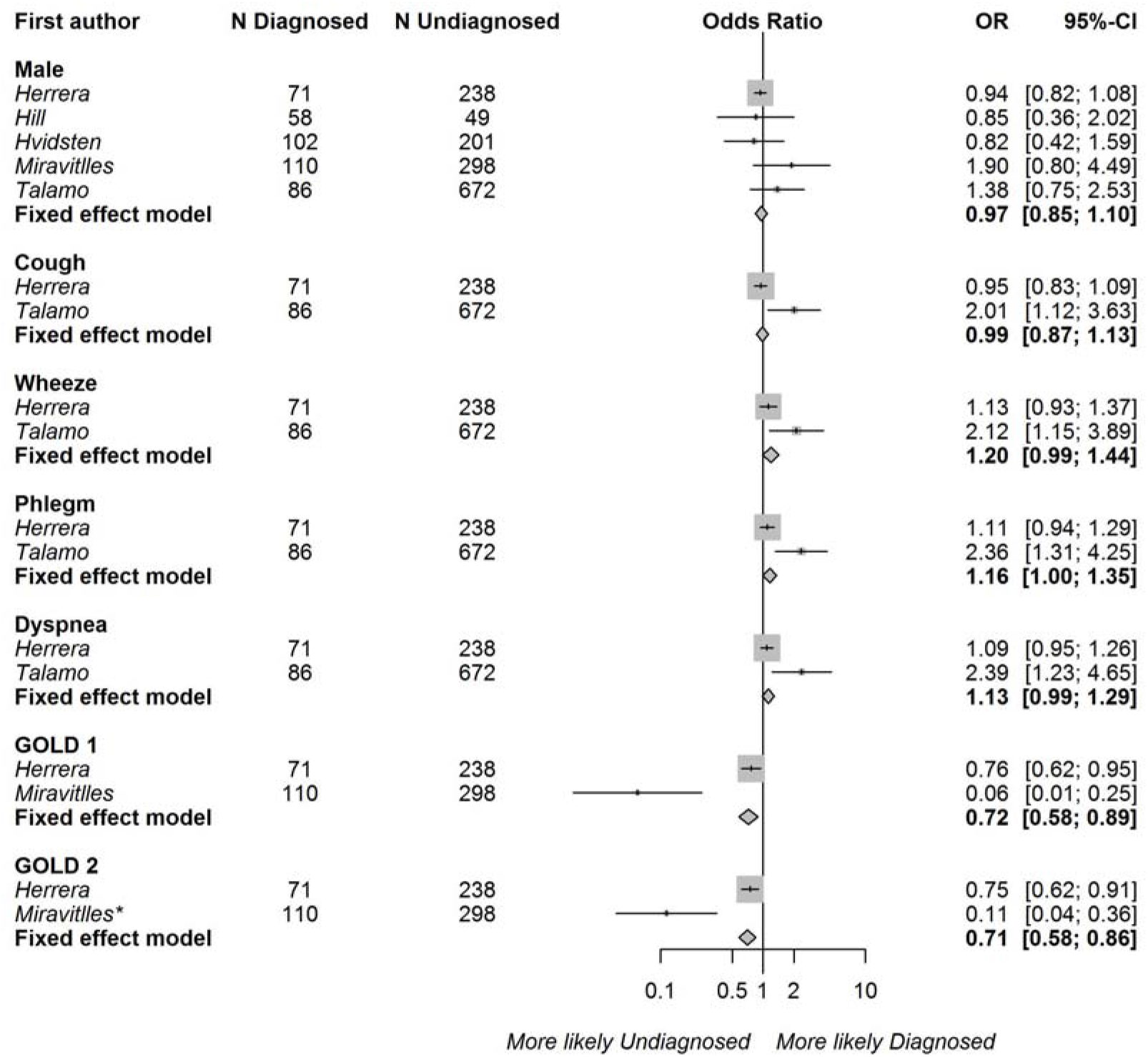
Associations between risk factors and the odds of receiving a COPD diagnosis using the regression coefficients from studies with multivariable regression modelling† and persistent airflow limitation defined as post-bronchodilator FEV_1_/FVC<0·7. The reference categories were female, the absence of cough, wheeze, dyspnoea, phlegm, and GOLD grades 3 and 4, respectively. Squares represent individual study estimates with the size of the square corresponding to their weight in the pooled estimate (represented with diamonds). *The reference category was changed from GOLD grade 1 to GOLD grades 3 and 4 by assuming a covariance of 0 between the dummy variables representing GOLD grades 1 and 2. Regression models were adjusted for age (*Herrera, Hill, Hvidsten, Miravitlles, Talamo)*, sex (*Herrera, Hill, Hvidsten, Miravitlles, Talamo),* ethnicity *(Herrera, Talamo)*, body mass index *(Herrera, Hvidsten)*, education (*Herrera, Hvidsten, Miravitlles, Talamo)*, income *(Hvidsten)*, employment *(Talamo)*, risk factor to dust *(Herrera)*, smoking *(Herrera, Hill, Hvidsten, Miravitlles, Talamo)*, respiratory symptoms, *(Herrera, Hill, Hvidsten, Talamo)*, self-rated health *(Hvidsten, Miravitlles)*, COPD severity *(Herrera, Miravitlles, Talamo)*, comorbidities *(Herrera, Hvidsten)*, prior health-care use (*Herrera, Hill)*, and exacerbations *(Herrera)*.

**Figure 5:**
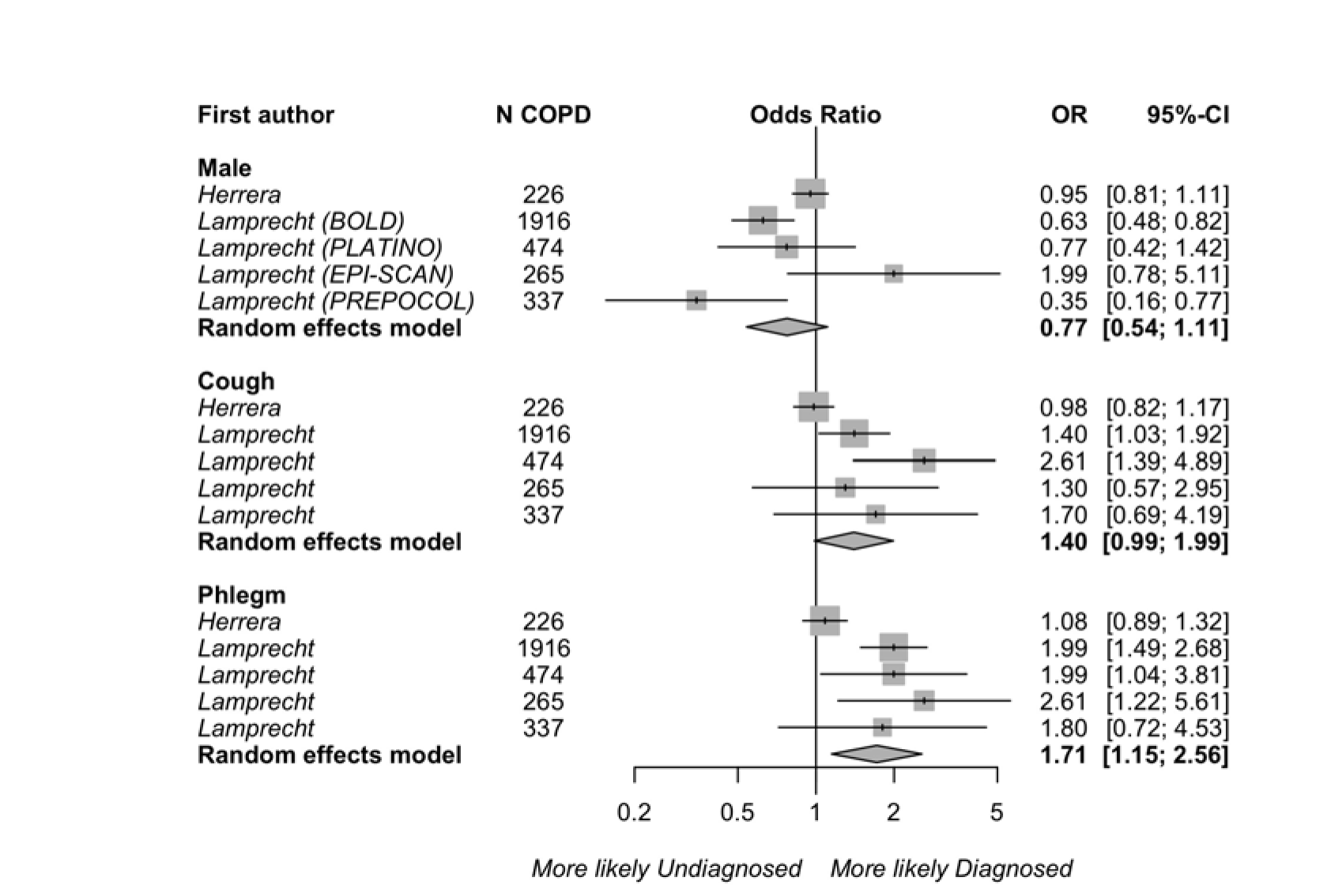
Associations between risk factors and the odds of receiving a COPD diagnosis using the regression coefficients from studies with multivariable regression modelling and persistent airflow limitation defined as post-bronchodilator FEV1/FVC<LLN. The reference categories were female, and the absence of cough and phlegm, respectively. The results for each dataset (BOLD, PLATINO, EPI-SCAN, PREPOCOL) analysed in Lamprecht et al.^4^ were pooled separately. Squares represent individual study estimates with the size of the square corresponding to their weight in the pooled estimate (represented with diamonds). †*Herrera et al.*^23^ reported prevalence ratios from Poisson regression models. Regression models were adjusted for age (*Herrera, Lamprecht)*, sex *(Herrera, Lamprecht)*, ethnicity *(Herrera)*, body mass index *(Herrera)*, education *(Herrera, Lamprecht)*, risk factors to dust *(Herrera)*, smoking *(Herrera, Lamprecht)*, respiratory symptoms (*Herrera, Lamprecht)*, COPD severity *(Herrera, Lamprecht),* comorbidities *(Herrera)*, and prior health-care use *(Herrera, Lamprecht)*.

Three risk factors were pooled in our assessment of studies using adjusted analysis based on the LLN definition of persistent airflow limitation. This analysis indicated a more strongly positive association between the presence of phlegm and being diagnosed with COPD (OR 1·71, 95% CI 1·15-2·56), although there was heterogeneity between datasets (I^2^ 75·2%). Patient sex and the presence of cough had no independent effect.

## Discussion

The presence of respiratory symptoms and GOLD 3 or 4 disease severity was strongly associated with a prior diagnosis of COPD among individuals with persistent airflow limitation on spirometry. These findings were relatively consistent across analysis methods and alternate definitions of persistent airflow limitation. Greater disease severity was the most important characteristic of diagnosed patients in two out of three pooled analyses in which spirometry was performed. In particular, patients with mild or moderate COPD (as measured by GOLD grades) were 78% less likely to have received a diagnosis than patients with severe or very severe COPD in the unadjusted analysis (based on contingency tables), and mean percent predicted FEV1 was 13% lower in diagnosed than undiagnosed patients. Disease severity was also the only risk factor that was associated with a diagnosis in both the unadjusted and adjusted (based on regression modelling) analyses. In the adjusted analysis, patients with moderate COPD were 29% less likely to have received a diagnosis than patients with severe or very severe COPD. Respiratory symptoms were another group of risk factors that were correlated with a COPD diagnosis. Among respiratory symptoms, the presence of dyspnoea was the most strongly associated with a previous diagnosis in the unadjusted analysis. Patients with ‘diagnosed’ COPD scored 0·52 points higher on the mMRC dyspnoea scale. However, there was only one study^25^ in which the mean score on the mMRC scale could have been used to distinguish undiagnosed from diagnosed patients using commonly accepted criteria (‘more dyspnoea’ if mMRC score ≥2 v. ‘less dyspnoea’ if mMRC score <2).^1^ Following dyspnoea, the presence of wheeze, and phlegm was also strongly associated with ‘diagnosed’ COPD in the unadjusted analysis. However, in the adjusted analysis, phlegm was the only symptom that was independently associated with having received a diagnosis, and this association was weaker than the unadjusted one. Interestingly, the presence of coughing was not well associated with a previous diagnosis in any of the pooled analyses. Overall, aside from the attenuated results in the adjusted analysis (discussed in detail below), our findings suggest a strong association between the presence of dyspnoea, phlegm, or wheeze and a COPD diagnosis. In addition to patients with respiratory symptoms being more likely to seek care, current guidelines now consider the presence of symptoms as part of the criteria for diagnosing COPD among patients with persistent airflow limitation^1^.

Patient characteristics such as sex and age were not associated with an increased likelihood of having received a diagnosis in any of the pooled analyses. There was some indication that patients with ‘diagnosed’ COPD had a greater pack-year smoking history, although current smoking status and smoking history were not statistically significant when they were assessed as the presence of former smoking and never smoking. The effects of risk factors on the likelihood of being diagnosed were weaker in the adjusted analyses than in the unadjusted analyses. The adjusted analyses were based on pooled coefficients from regression modelling. Although the inclusion of covariates is expected to reduce the effects sizes compared to odds ratios derived from contingency tables (as in the unadjusted analysis), one study in the adjusted analysis^23^ had unusual results that received disproportionate weighting. In contrast to all other studies in this review, Herrera et al.^23^ found that respiratory symptoms were not associated with the likelihood of having received a diagnosis of COPD. In the adjusted analysis, these results were pooled with one other study^17^, which found that the presence of respiratory symptoms strongly impacted the likelihood of receiving a diagnosis. This discrepancy between studies may be due to differences in the population that was sampled (primary care clinic^23^ versus general population^17^). In general, studies in clinic settings might have observed smaller differences between undiagnosed and diagnosed patients because they sampled from a subset of patients that were prompted to seek care because of a symptom burden.

Our systematic review has several strengths. First, we used data from a total of 16 articles in the meta-analysis, and these articles were mostly population-based studies that scored high in quality. Second, there were a robust number of studies for many risk factors; patient sex was assessed in 10 studies in total, followed by disease severity in 9 studies, and respiratory symptoms and smoking history in 8 studies each. The methods used to measure disease severity, respiratory symptoms, and smoking history were relatively consistent across studies, which facilitated pooling of their findings. Lastly, we conducted several pooled analyses to assess the sensitivity of our findings to alternate definitions of COPD (fixed ratio and LLN) and analysis methods (unadjusted and adjusted analyses). Except for one study,^23^ our findings were consistent.

However, our systematic review also has several limitations. First, half of the pooled samples were based on data from three large prevalence studies (EPI-SCAN, PLATINO, and BOLD). This resulted in overrepresentation of patients in Spain and Latin America; differences in patient and physician behaviour and health-care services use can result in findings that vary across settings. Second, although the total number of studies for each risk factor was robust, the number of studies assessing each risk factor within pooled analyses tended to be small. This was partly because separate articles using the same dataset could not be combined in our pooled analyses. The number of studies used in the unadjusted analysis of respiratory symptoms and the adjusted analysis using the LLN definition of COPD was reduced as a result. Third, with the exception of dyspnoea, all other respiratory symptoms in the pooled analyses were measured as binary variables (either present or absent). Given our finding that symptoms are characteristic of a COPD diagnosis, a more nuanced assessment of their severity might result in an even greater ability to distinguish between undiagnosed and diagnosed patients. In addition, because respiratory symptoms were self-reported in all studies, reporting bias might have exaggerated the difference in symptoms between the undiagnosed and diagnosed groups. The findings from this systematic review have important implications for research and policy around COPD diagnosis, for example, in estimating the return on investment in screening and case detection strategies for COPD. The true burden of COPD is the sum of the disease burden in diagnosed and undiagnosed patients, and our results indicate that undiagnosed patients generally have milder disease and therefore a lower disease burden.

On one hand, this indicates that strategies aiming to reduce the underdiagnosis problem are unlikely to result in immediate and dramatic improvements in patient-related outcomes such as symptom burden. On the other hand, the gap in disease severity and symptom burden between diagnosed and undiagnosed patients indicates a delay in COPD diagnosis among patients that have already developed symptoms. Given the potential for disease modification at early stages of COPD, reducing this delay could be associated with substantial improvement in long-term patient outcomes and a reduction in mortality and costs.

## Acknowledgements

This study was funded by the Canadian Institutes of Health Research (application number 142238).

## Appendix

**Text A1:**
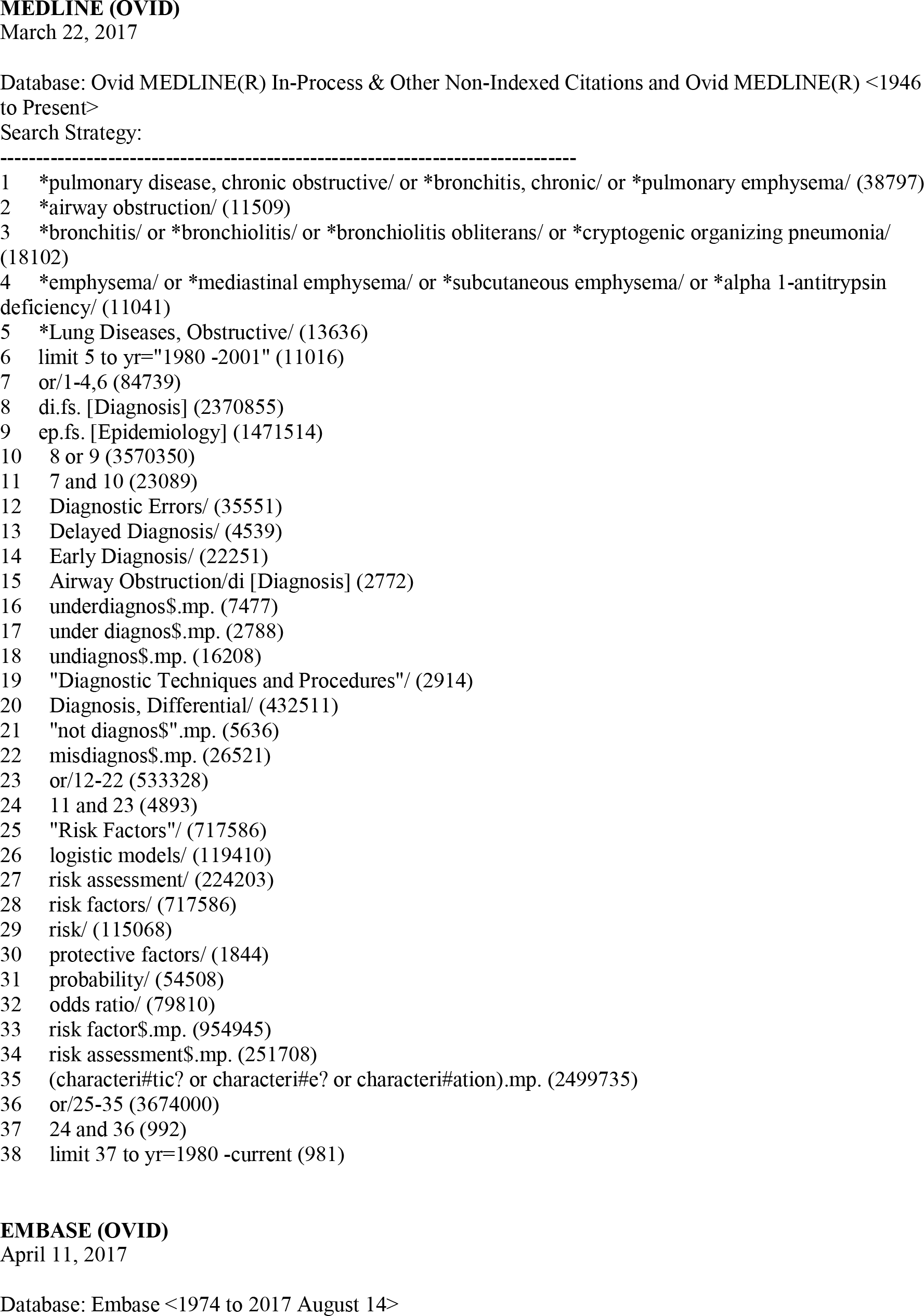
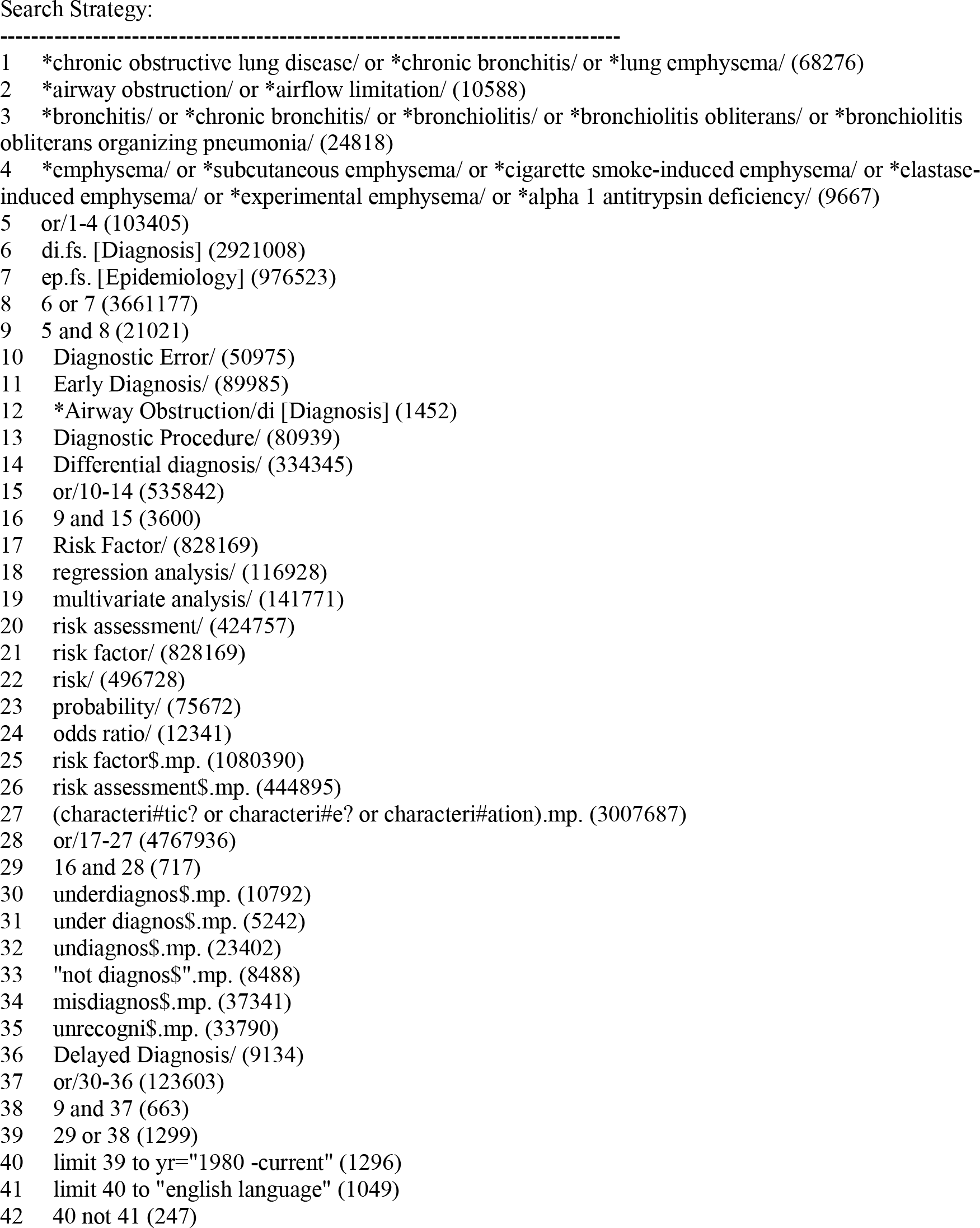
Search strategy

**Figure A1:**
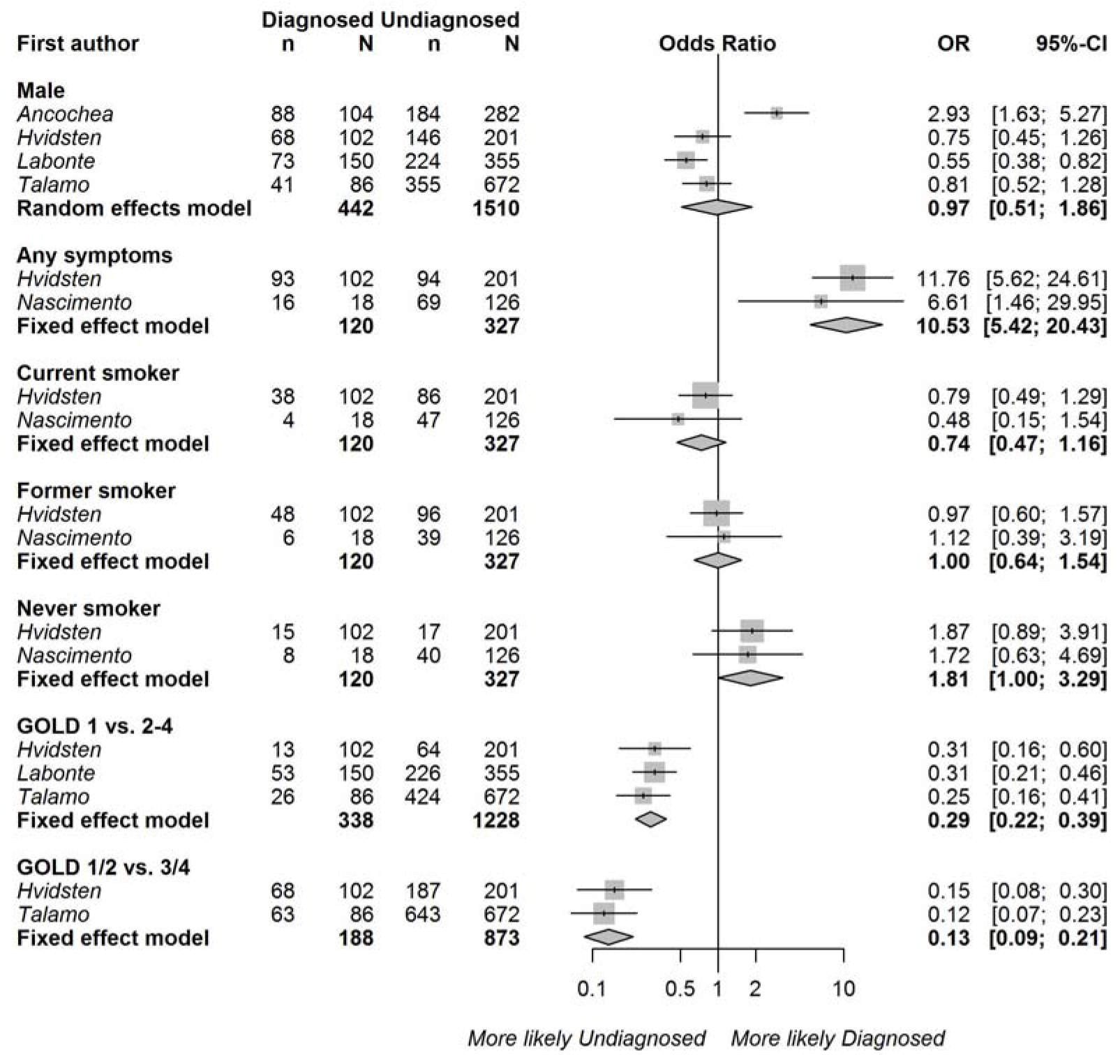
Associations between diagnosed (v. ‘undiagnosed’) COPD and sex, the presence of any respiratory symptoms, smoking status, smoking history, and COPD severity based on the contingency tables of studies using random sampling of the general population. Persistent airflow limitation was defined as post-bronchodilator FEV_1_/FVC<0·7. Squares represent individual study estimates with the size of the square corresponding to their weight in the pooled estimate (represented with diamonds).

**Figure A2:**
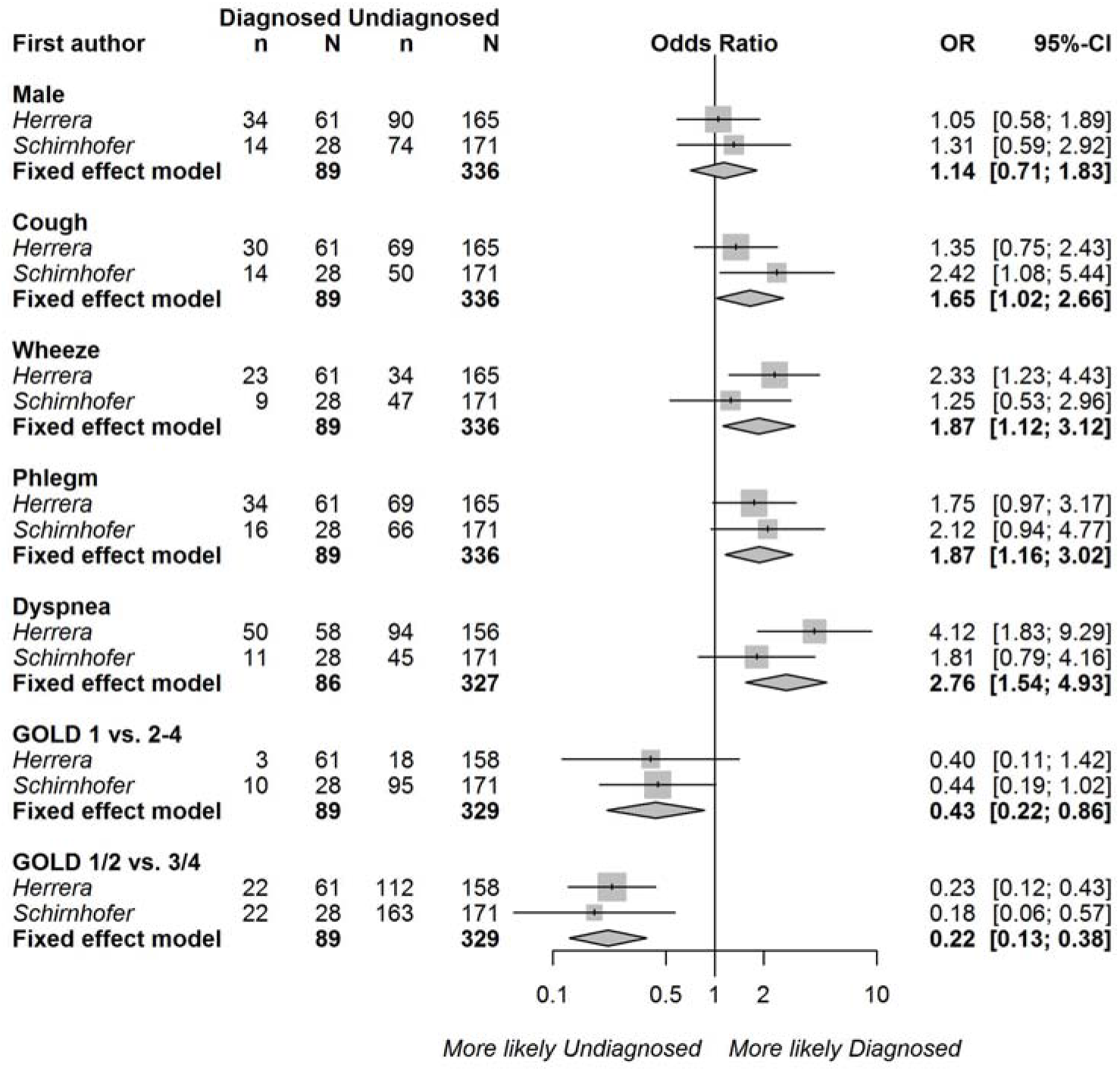
Associations between diagnosed (v. ‘undiagnosed’) COPD and sex, the presence of cough, wheeze, phlegm, dyspnoea, and COPD severity based on contingency tables. Persistent airflow limitation was defined as post-bronchodilator FEV1/FVC< lower limit of normal (LLN). Squares represent individual study estimates with the size of the square corresponding to their weight in the pooled estimate (represented with diamonds).

